# Do substrate roughness and gap distance impact gap-bridging strategies in arboreal chameleons

**DOI:** 10.1101/2020.08.21.260596

**Authors:** Allison M. Luger, Vermeylen Vincent, Herrel Anthony, Adriaens Dominique

**Affiliations:** Ghent University, Evolutionary Morphology of Vertebrates – Ghent, Belgium; C.N.R.S./M.N.H.N. Département Adaptations du Vivant, Bâtiment d’Anatomie Comparée – Paris, France

**Author notes:** **Cite as:** Luger, A.M., Vermeylen, V., Herrel, A. and Adriaens, D. (2020) Do substrate roughness and gap distance impact gap-bridging strategies in arboreal chameleons? bioRxiv, 2020.08.21.260596, ver. 3 peer-reviewed and recommended by PCI Zoology. doi: https://doi.org/10.1101/2020.08.21.260596. This article has been peer-reviewed and recommended by *Peer Community in Zoology* doi: https://doi.org/10.24072/pci.zool.100005.

**Keywords:** Chameleons, Prehensility, Gap Bridging, Substrate Type

## Abstract

Chameleons are well-equipped for an arboreal lifestyle, having ‘zygodactylous’ hands and feet as well as a fully prehensile tail. However, to what degree tail use is preferred over autopod prehension has been largely neglected. Using an indoor experimental set-up, where chameleons had to cross gaps of varying distances, we tested the effect of substrate diameter and roughness on tail use in *Chamaeleo calyptratus*. Our results show that when crossing greater distances, *C. calyptratus* is more likely to use its tail for additional stability. The animals were able to cross greater distances (up to 1 75 times the shoulder-hip length) on perches with a rougher surface. We saw that depending on the distance of the gap, chameleons would change how they use their prehensile tails when crossing. With shorter gaps the tails either do not touch, or only touch the perch without coiling around it. With larger distances the tails are fully coiled around the perch, and with the largest distances additionally they reposition the hind legs, shifting them towards the end of the perch. Males were able to cross relatively greater distances than females, likely due to their larger size and strength.

## Introduction

Arboreal habitats consist of complex three-dimensional structures with discontinuities, thus leaving gaps between solid surfaces (Cartmill, 1974). Animals living in such arboreal environments are challenged with the task of crossing these gaps in order to create a more direct path through the canopy, avoiding long and possibly dangerous and costly detours to reach their destination. When crossing, an animal will either have to reach for the other branch, leaving a portion of the body unsupported, or alternatively leap, glide, or fly to reach the other side of the gap. These strategies all impose different mechanical demands on the locomotor system of the animal. While most studies have focused on the horizontal movement over branches, there have been some studies that have focused on the movement between branches, such as in arboreal snakes (Hoefer et al., 2013; Byrnes et al., 2012) or primates (Thorpe et al., 2009). An extensive study on the various methods of gap crossing was performed by Graham et al. (2020), though this did not include the use of prehensile tails. Peterson (1984) studied the locomotion of chameleons, focusing on the adaptations of the limbs and tail. Chameleons, which are not known for their leaping abilities, move between perches by extending their front limbs, relying on their hind limbs and tail for support. The pectoral girdle is not fixed relative to the body wall allowing it to slide and rotate further in the sagittal plane, giving it a greater reach allowing chameleons to cross larger distances compared to other arboreal squamates such as anoles (Peterson, 1984).

Several morphological and functional adaptations have evolved in chameleons to accommodate an arboreal lifestyle including ‘zygodactylous’ hands and feet and a fully prehensile tail (Gans, 1967). Their ‘zygodactylous’ hands and feet form grasping appendages that can hold onto narrow branches, while their tail has modifications both in the musculature and the morphology of the caudal vertebrae (Ali, 1948; Zippel et al., 1999; Bergmann et al., 2003; Luger et al., 2020). These characteristics serve multiple purposes for arboreal chameleons and are not only restricted to movement on narrow perches in the forest canopy, climbing, or bridging gaps but are also important during social behavior. Indeed, when males encounter each other, this often results in fighting during which a male will attempt to toss the other from the shared branch (Measey et al., 2009).

Long tails are often thought to be adaptations for an arboreal lifestyle in chameleons (Bickel and Losos, 2002). Prehensile-tailed arboreal chameleons have longer tails with more vertebrae, compared to terrestrial species. This allows them to coil their tail multiple times around a perch, thereby increasing the contact area and thus friction. This results in an improved tail gripping performance compared to non-prehensile-tailed chameleon species (Herrel et al., 2012; Luger et al., 2020). While the effect of hand/feet size and tail length on gripping abilities has been studied, little is known, however, on how chameleons use their prehensile tail on different substrates or perches of different diameters. The use of the tail to increase gripping performance can be expected when bridging large gaps between branches, or when having to grip onto smooth, low-friction substrates. Therefore, we studied how chameleons change the way they use their prehensile tail when confronted with different conditions including gaps of different sizes, different substrate diameters, or different substrate roughness. In addition, we examined whether this behavior differs between sexes.

*Chamaeleo calyptratus* (Duméril and Bibron, 1851), a large-sized arboreal chameleon, was used for this study. We first quantified whether chameleons changed the use of their tail when substrate roughness, perch diameter, and gap distance were modified using an indoor experimental set-up. First, we predicted an increase in tail use on smoother substrates as this decreases the grasping ability of the hands and feet.

Previous studies by Spinner et al. (2014) and Khannoon et al. (2014) showed a relationship between substrate roughness and friction. Second, we predicted that when bridging greater gaps, animals would suspend their body while gripping the perch with their hind legs and tail. When crossing greater gaps, the center of mass of the animal thus being suspended further from the points of attachment in the feet and tail. Third, we predicted that animals more often use their tail as an anchor as gap distance increases. Our fourth prediction was that when on perches with a rougher surface, chameleons should be able to cross greater gap distances compared to smoother surfaces. Finally, many chameleons, including *C. calyptratus*, are sexually dimorphic in body size, tail length and hand and feet span (Bickel and Losos, 2002). In dwarf chameleons of the species *Bradypodion* males also have larger hands and feet and longer tails resulting in a higher gripping force compared to females (Herrel et al., 2011; Da Silva et al., 2014). Consequently, we predicted differences in how chameleons use their prehensile tail based on sex-related size variation. We specifically predict that males are able to cross greater distances as their increased hand and foot length gives them a higher gripping strength, increasing friction and stability, at least on larger perches. We also predicted males to make more successful crossings without using their tail due to their expected higher grip strength.

## Methods

### Animals

*Chamaeleo calyptratus* specimens (*N* = 9; five males and four females) were obtained via a private breeder in Paris, France at 10 weeks of age. The animals were at least one year old and sexually mature at the start of the experiments. We chose to use *C. calyptratus* because of their availability, easy husbandry, and resilience under stressful conditions. The animals were kept in wire cages of 90 × 45 × 46 cm and fed *ad libitum* on a diet of vegetables (lettuce) and crickets dusted with mineral and vitamin supplements. Each cage was provided with an UV HID-Lamp, creating a thermal gradient within the environment. Room temperature was kept constant at 26°C, which is consistent with their natural ambient temperature. The humidity was kept between 20-50% and the animals were kept on a day/night cycle of 13/11h. The animals were checked weekly for any injuries or indications of disease, malnutrition or dehydration. Measurements of the animals can be found in Suppl. Table 1. According to the Belgian legislation on experimental animals, the chameleons were checked by a veterinarian every three months. The experimental design was approved by the Ethics Committee for Animal Experimentation of Ghent University (Faculty of Sciences) (application 2017-028).

### Perch crossing experiment

Videos were recorded using a JVC HD Everio camera. The experiments were performed in the same room in which the animals were housed. According to Andrews (2008) the selected body temperature of *C. calyptratus* is suggested to be at 30.4°C, as such the chameleons were tested only slightly below their selected body temperatures. Filming occurred between March 2018 and May 2019. Trials with the same set-up (material, perch diameter, etc.) were performed subsequently, after which the set-up was adjusted and the next trial commenced after a break of 30 minutes. When animals appeared to be tired or otherwise unwilling to cooperate, the session was terminated and continued on a later date. At least one day of rest was given between each filming session.

Two plastic holders (customized and 3D printed) in which the perches were inserted, were fitted on a vertical rod (Figure 1). The holders were manually adjusted to alter the gap distance (measured using a handheld ruler). Perches of three materials with a different degree of roughness were used (from smoothest to roughest): PVC, wood, and 3M sandpaper with a roughness factor of P100 (attached to a wooden perch). Of each material, two different perch diameters were used: a narrow one of 9 mm and a broad one of 25 mm. The diameter of the perches was chosen based on the range of commercially available rods. Gap distances between the two perches were standardized at 0.5, 1, 1.25, 1.5, 1.75, and 2 times the shoulder-hip length, averaged by sex. An average per sex was used instead of per individual, as the individual variation was low. For each individual we measured the shoulder-hip length using digital calipers and took the average per sex for the experiments. For females, with an average shoulder-hip length of 110 mm, the gap distances were set at 55, 110, 137.5, 165, 192.5, and 220 mm. For males, with an average shoulder-hip length of 130 mm, this was 65, 13, 162.5, 195, 227.5, and 260 mm. A total of 175 trials in which a successful crossing was made were recorded for the wooden perches (75 for females, 100 for males), 13 for the PVC perches (2 for females, 11 for males), and 139 for the sandpaper perches (64 for females, 75 for males).

**Figure 1:**
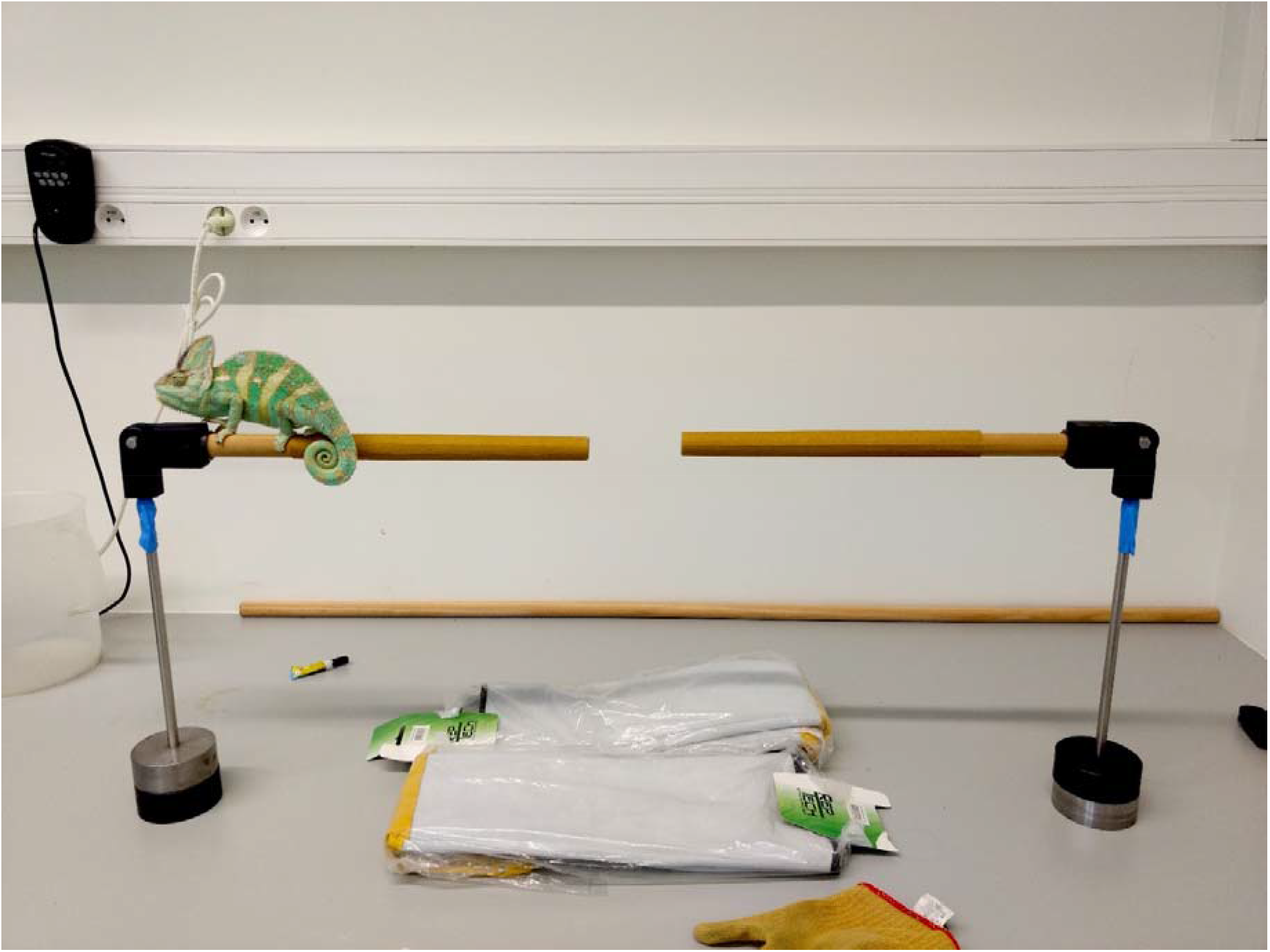
Experimental set-up used. The set-up consists of two holders in which perches of various sizes could be inserted. The holders could be moved manually adjusting the distance between perches.

### Tail use analysis

The video data was analyzed using VLC Mediaplayer. To examine the relationship between prehensile tail use and perch material and diameter, we counted the number of times the chameleon used or did not use its tail to assist it to bridge the gap. The animal was considered to be using its tail if at any point during the crossing of the gap, it would grasp the perch with its tail, coiling it around it. We plotted the number of times the tail was either used or not used against substrate type and diameter. We then calculated the percentage of times an animal managed to cross the predetermined gap distance per set-up with or without the tail (Suppl. Table 1). We focused on the various tail grasping strategies a chameleon applied when successfully crossing the gap (unsuccessful crossing was thus not considered). This study is thus mainly of a descriptive nature, focusing on qualitative traits in tail use.

## Results

### Tail use

The number of times the animals used their tail is listed in Suppl. Table 2 and plotted against perch thickness and material (Fig. 2 for females, Fig. 3 for males). For the smallest gap (0.5 times shoulder-hip length), the tail was most often not used (average for both sexes: 72% of the trials). When on broad perches, animals use their tail more often for both wood and sandpaper. For greater distances, tail use frequency increased (on average in 87% of the trials for 1, 1.25 and 1.5 times the shoulder-hip length). Only for one trial with a gap equal to 1.5 times the shoulder-hip length a successful crossing was made without using the tail. Otherwise all other successful crossings using wood and sandpaper for gaps equal to 1.5 and 1.75 times the shoulder-hip length involved the use of the tail. Sandpaper was the only material for which the chameleons managed to cross a gap distance of 1.75 times their shoulder-hip length. At twice the shoulder-hip length, no successful crossings were observed, irrespective of the material. For PVC, crossings were less successful than with the other materials. Many trials were conducted with the PVC perches, but in the end only a total of 13 successful crossings were observed. On the broad PVC perch chameleons were able to bridge gaps of 0.5 times the shoulder-hip length only. On narrow perches some successes were observed at gaps equal to the shoulder-hip length, though only for males.

**Figure 2:**
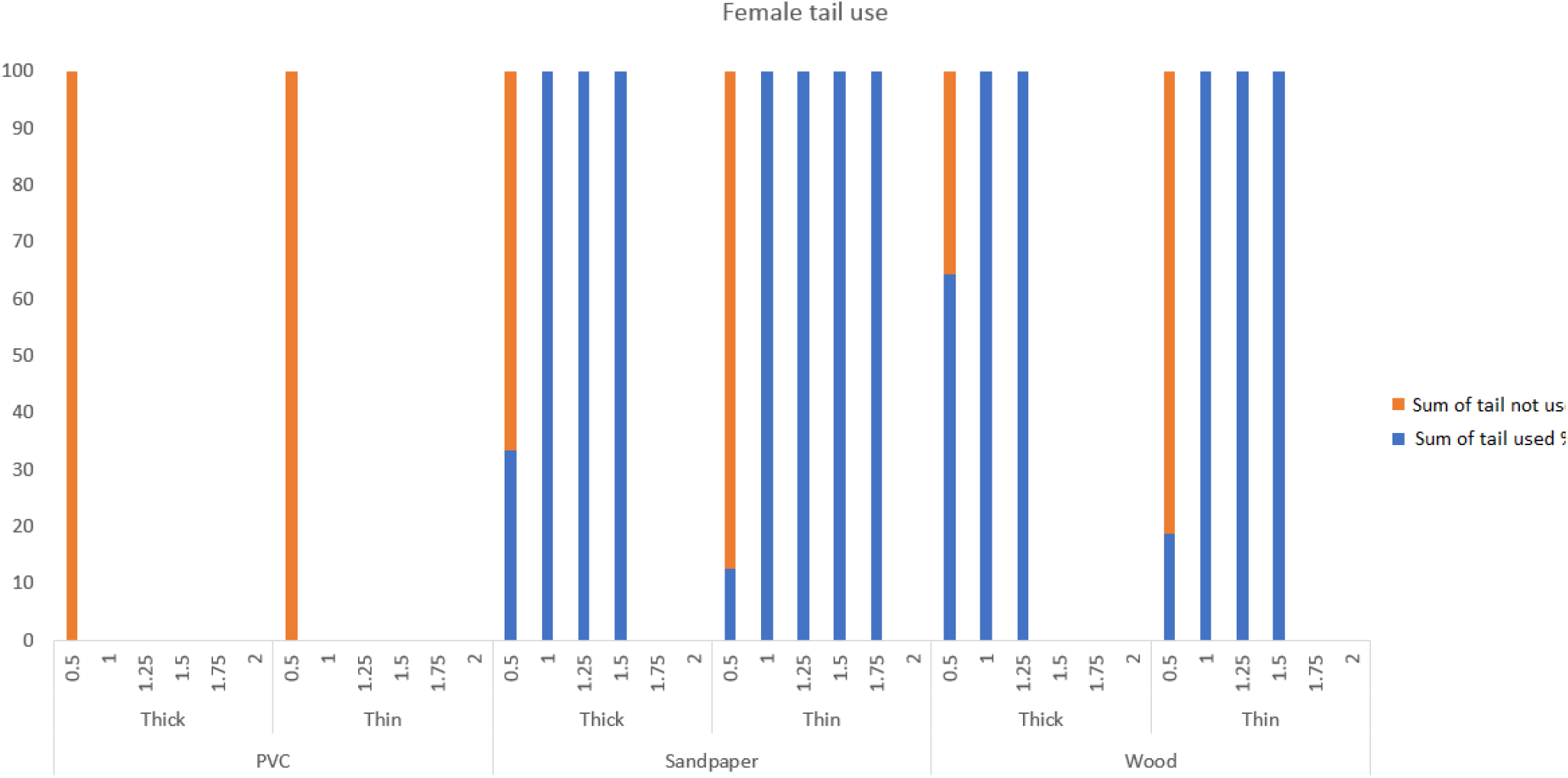
Frequency of tail use plotted against perch thickness and material in female *Chamaeleo calyptratus*, the absolute number of successful trials can be found in Suppl. Table 2. Percentages given are the number of times a tail was used or not for each of the three different materials (PVC, sandpaper and wood) and the two different perch diameters (narrow and broad). Distances without any data indicate that no successful crossings were made for the trials with that perch diameter and material.

**Figure 3:**
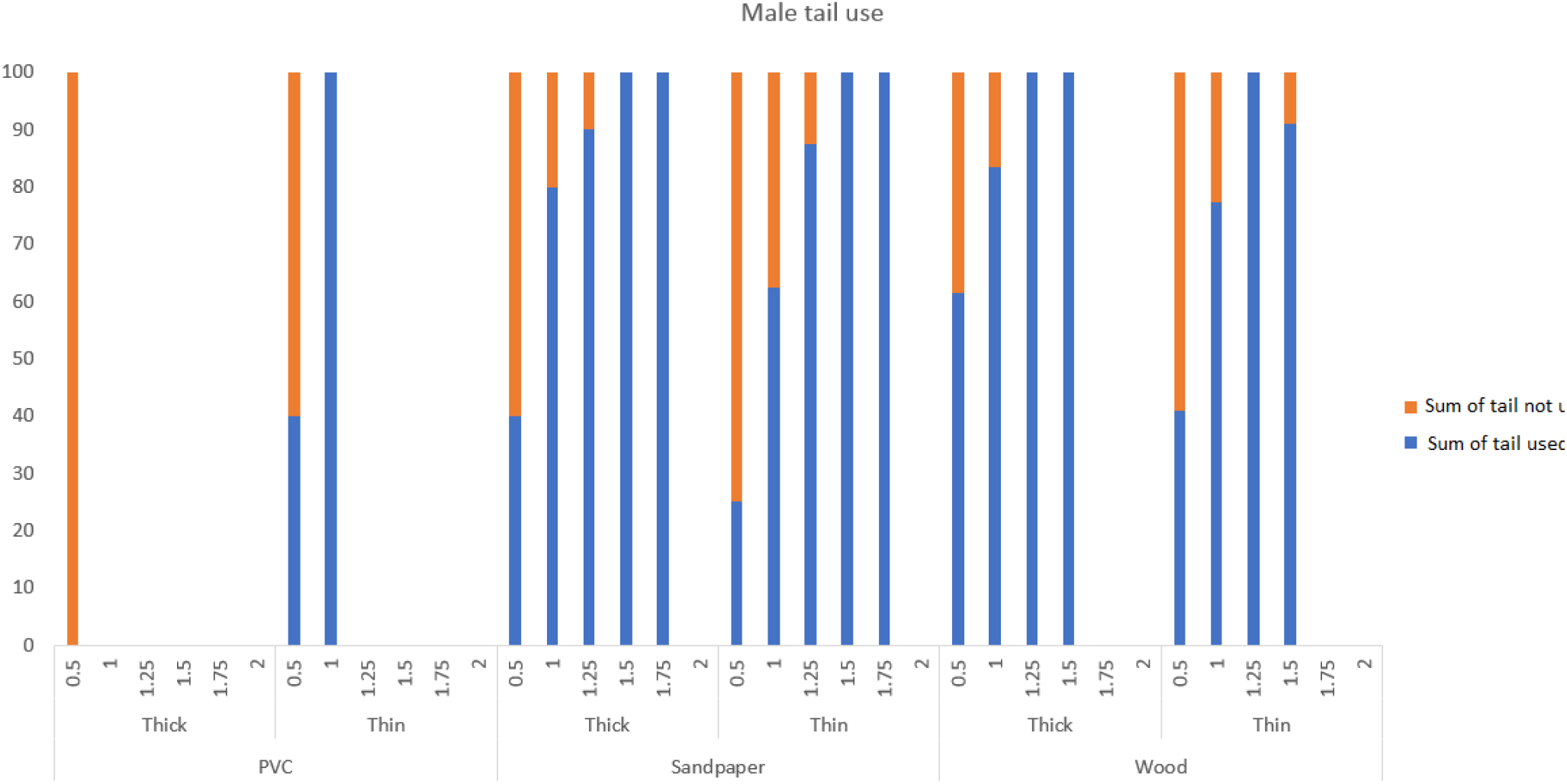
Frequency of tail use plotted against perch thickness and material in male *Chamaeleo calyptratus*, the absolute number of successful trials can be found in Suppl. Table 2. Percentages given are the number of times a tail was used or not for each of the three different materials (PVC, sandpaper and wood) and the two different perch diameters (narrow and broad). Distances without any data indicate that no successful crossings were made for the trials with that perch diameter and material.

Males made less use of their tail to successfully cross gaps compared to females. Females only occasionally crossed gaps of 0.5 times their shoulder-hip length without using their tail. In all other trials they coiled their tail around the perch. Females were also less capable of crossing wider gaps than males, with or without tail. Only on the narrow sandpaper perches did females cross a distance of 1.5 times their shoulder-hip length. Males would still attempt crossing distances of 1 and 1.25 times the shoulder-hip length without using their tail; for example, 28% of the sandpaper trials at shoulder-hip length did not involve the use of the tail and 11% for a distance of 1.25 times the shoulder-hip length. On wood perches, males did not use their tail in 20% of the trials with gap distances equal to their shoulder-hip length. The tail was always used, however, to bridge gaps of 1.25 times the shoulder-hip distance on wood perches. One male chameleon succeeded to cross the distance of 1.5 shoulder-hip distance on a narrow wooden perch without using its tail. For all other trials with gap distances of 1.5 and 1.75 times the shoulder-hip length the tail was used.

### Gap crossing strategies

For the shortest distances, two different strategies were observed. Only the smallest gap distance (0.5 times shoulder-hip length) could be traversed easily regardless of perch material and diameter and most often without using the tail. For such short distances, the chameleon would not stop and pause to coil their tail around the perch, but without showing any anticipatory behavior would continue walking using their regular gait. A second strategy for shorter gaps is observed when using their tail for gaps equal to 0.5 and 1 time shoulder-hip length. The chameleon would first grab the next perch with its hands, coil the tail around the perch, and then proceed to move the hind legs to the next perch, before letting its tail go. Often the tail would not even be fully coiled, only wrapping it halfway along the underside of the perch leaving the distal end suspended (Fig. 4). We refer to this strategy as the “better-safe-than-sorry”-strategy, as chameleons had proven to be able to safely cross these distances without using their tail.

**Figure 4:**
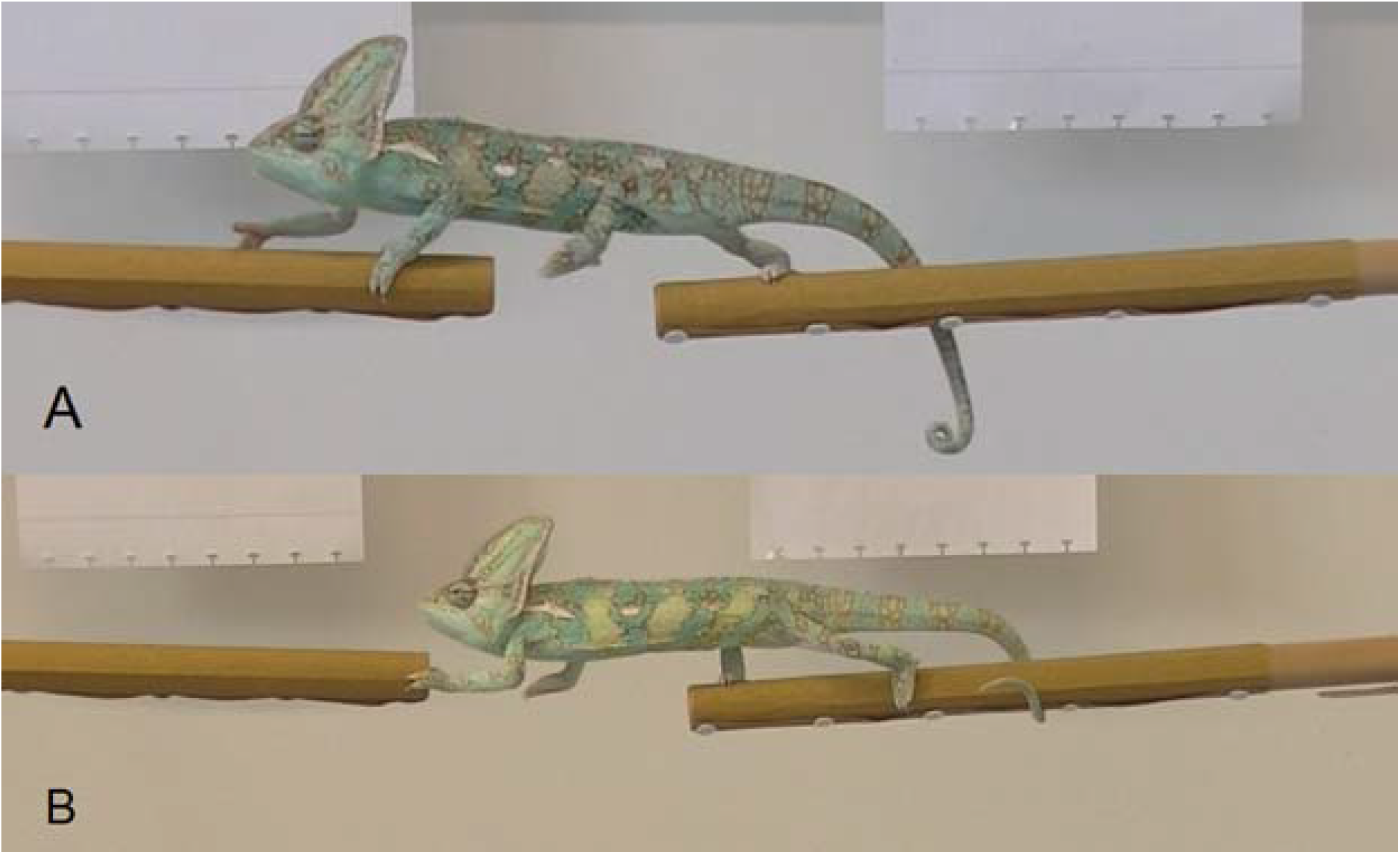
Tail use strategies for short distances. For the shortest distances (0.5 or 1 times the shoulder-hip length) the chameleon *Chamaeleo calyptratus* would employ two different strategies: one with a regular locomotion taking larger steps to cross the gap (A) or the “better-safe-than-sorry”-strategy coiling the tail half around the perch when crossing (B).

With gap distances above 1.25 times shoulder-hip length, we observed a different strategy. They would start coiling their tail around the perch before grabbing the next perch with their hands. After their hands grabbed onto the next perch, they would slightly uncoil the tail, slide it further along the perch, and re-coil the tail at a new position closer to the gap. After they had repositioned the tail, they would move both hind legs at the same time across the gap, before releasing their tail (Fig. 5). For even greater distances, chameleons would release their tail before they could reach the next perch with their hind legs, leaving them suspended only holding on by their fore limbs (Fig. 6). In these instances, only with sandpaper and on the narrow perch would the chameleon have enough grip in its hands to pull itself up fully to the next perch. The animal would grab the next perch with its hands and the departing perch with its tail, leaving the feet suspended (Fig. 6A). With broad perches, they would often fall before even an attempt to cross could be made.

**Figure 5:**
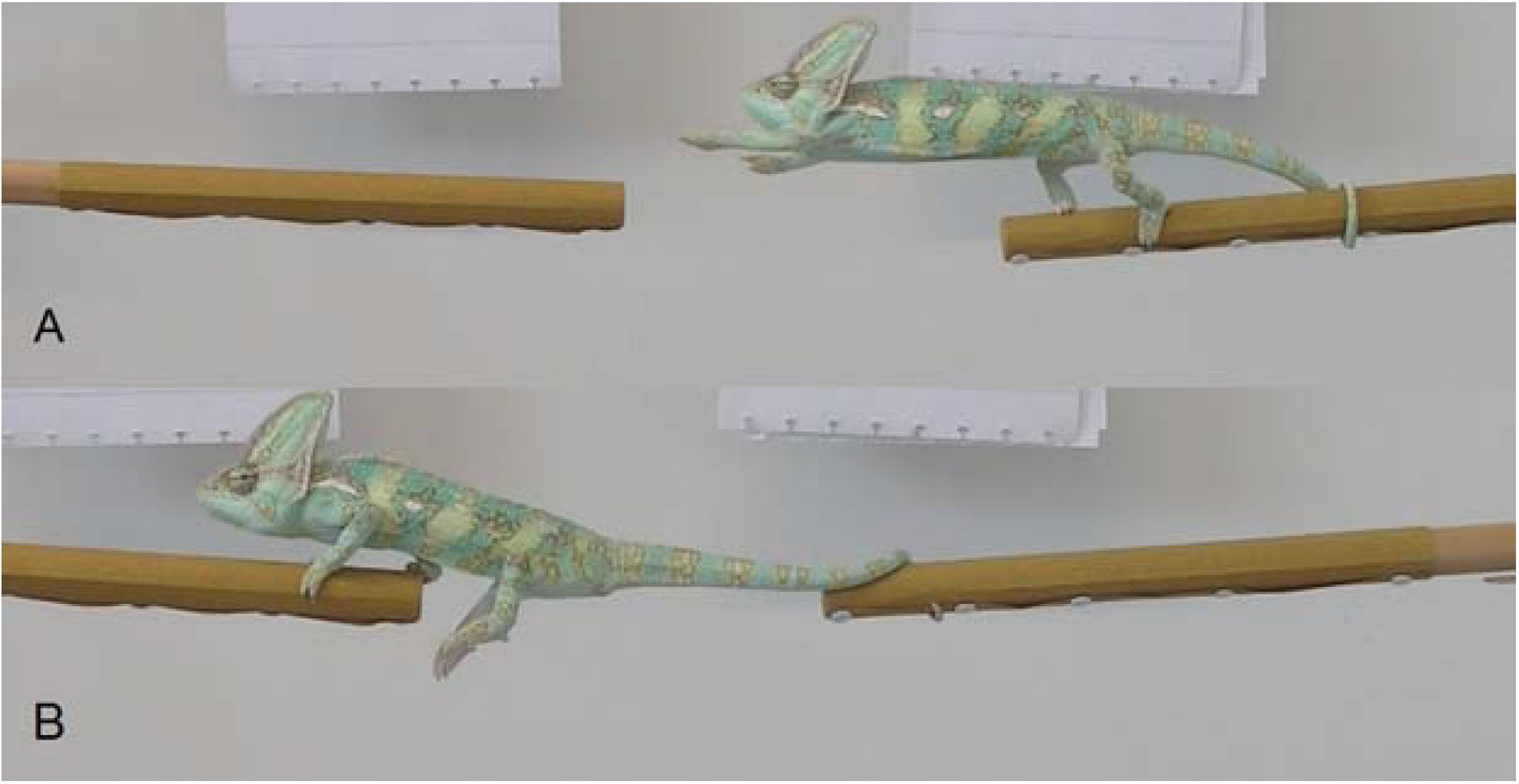
Tail use strategy used for crossing greater gap distances. For crossing a distance of >1.25 times its shoulder-hip length. The chameleon *Chamaeleo calyptratus* would coil its tail around the perch allowing it to grab onto the next one with its front legs (A). After that the animal would reposition its tail when holding onto the next perch, forming a bridge and moving both hind legs up to the next perch at the same time (B).

**Figure 6:**
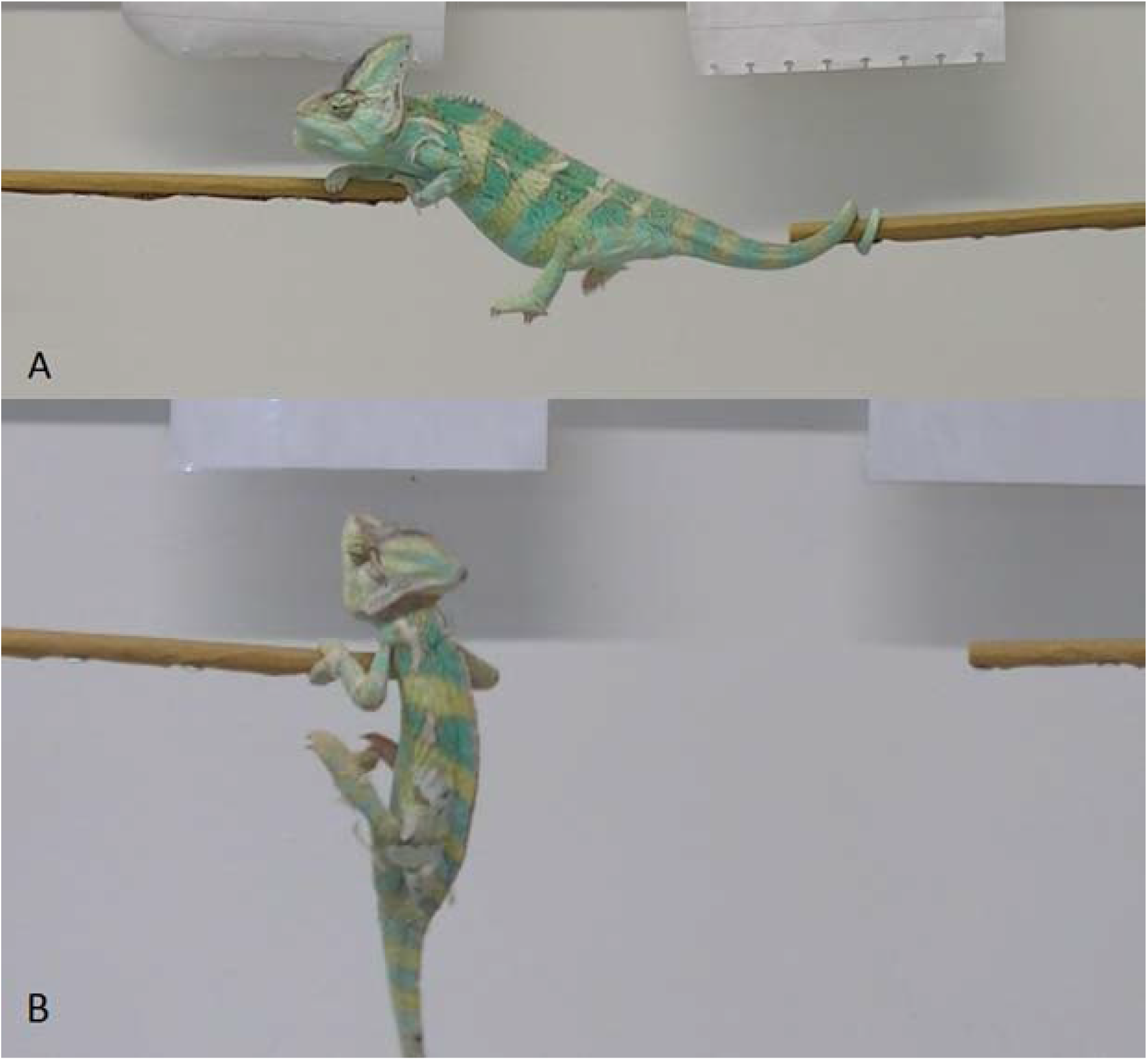
Second tail use strategy used for crossing greater gap distances. For crossing a distance of >1.5 times its shoulder-hip length. The chameleon *Chamaeleo calyptratus* would coil its tail around the perch allowing it to grab onto the next one with its front legs (A). Sometimes the distance would be too large to cross with the tail still holding on to the departing perch. In these instances. The animal would release the hind limbs and tail, holding only to the next perch with its hands having to pull up its entire body weight relying solely on the strength in its forelimbs.

## Discussion

### The effect of perch material and diameter in relation to crossing distance

Our results show that substrate smoothness and perch diameter have an effect on how *C. calyptratus* uses its tail. Moreover, the distance between perches (gap distance) impacts how and whether the tail is used. The grasping abilities of the hands, feet, and tail thus appear to be impacted by the amount of friction the animal has on the perch. The surface properties of the perch impact the friction with the hands, feet and tail of the chameleon. Our hypothesis that coarser substrates increase friction and thus allow chameleons to cross greater distances was supported by our data, which is consistent with the findings of Spinner et al. (2014) and Khannoon et al. (2014). Sandpaper allowed animals to cross the greatest distances (gaps up to 1.75 times their own shoulder-hip length), for both narrow and wide perches, whereas for the smoothest substrate (PVC), only short distances could be crossed successfully (0.5 and 1 times their shoulder-hip length; Figs. 2, 3). A gap of 1.75 times the shoulder-hip length seems to be the limit, as no crossings were observed for gaps of twice the shoulder-hip length, in neither males nor females. Our results also show that the greater the gap distance, the more often chameleons use their tails as an extra anchor point, coiling it around the perch while reaching with their arms for the next perch. In this case the hind legs, grasping the perch, act as a fulcrum while the tail, coiling around the perch behind, acts as an anchor line. In this way, the chameleon keeps its balance while shifting its center of gravity away from its foot contact point. As the crossing distance increased, the need to use their tail did as well. For wood and sandpaper perches, most of the trials involved coiling the tail; for distances of 1 time the shoulder hip length the tail is used in 88% of the trials, for 1.25 it is used 97% and for 1.5 times the shoulder-hip length it is used 86%.

Increasing the distance of the gaps creates a challenge for the chameleon, as both the amount of unsupported mass is increased as well as the length of the moment arm. While unable to leap, the extreme anterior excursion of the shoulder girdle and the humerus allows the chameleons to reach across distances greater than its own shoulder-hip length (Peterson, 1984). Crossing greater distances between broad perches appeared more difficult than between narrow ones (Suppl. Table 2). Herrel et al. (2011) showed that animals have a perch diameter preference related to the span of their hands and feet and perform better on perches that closely match the span of their hands and feet. Unlike the broad diameter, the narrow perches allow the chameleon to almost completely wrap their hands and feet around it. This often made the difference between a successful crossing or not, as for some greater distances the animal was not able to place the hind limbs on the next perch before releasing the tail from the departing perch. In these instances, the chameleon has to rely only on the strength in their forelimbs to pull itself up (Fig. 6). While we have only been studying the chameleon species, *C. calyptratus*, we can assume that other prehensile-tailed arboreal chameleon species might have similar strategies, based on their similar morphology and lifestyle. This hypothesis serves as a good starting point for future studies. Other chameleon species that have a more terrestrial lifestyle, or live in the lower shrubbery, might hold different strategies in how they utilize their tails when crossing, and could also be used for future research.

### Gap bridging strategies

A distance of 0.5 times shoulder-hip length appears to be slightly greater than the regular distance of a single step during a normal gait, which is in line with the stance length of *C. calyptratus* found by Fischer et al. (2010), but not long enough to require a different strategy. During regular walking, the chameleon tail is usually not engaged (see Peterson, 1984). Adding their tail clearly is not always necessary for these short gaps, as at all times at least one hand and foot are in contact with the substrate when crossing. For a gap equal to the shoulder-hip length, chameleons contact the perch with their tail, but without coiling it around the perch (Fig. 4). This likely helps them to maintain balance without having to slow down. Indeed, when the chameleon fully coils the tail around the perch, they pause. Using this “better-safe-than-sorry”-strategy for the shorter distances occurs with no, or only a very short, pause. Chameleons likely rely on the substrate being coarse to generate friction as they continue to use these strategies when confronted with a smoother substrate and despite the fact that this can cause them to fall while crossing. Chameleons do not appear to adapt their crossing strategy in response to the substrate roughness as they will make an attempt and simply tend to fall more often. The same applies for the broad perches where falling was quite common when crossing greater distances (Fig. 6). With smoother substrates and broader perches, chameleons appear not to be able to generate enough force to lift themselves onto the next perch. In some cases when an attempt to cross a gap was started without using the tail, they would reconsider halfway during the attempt and coil it while crossing. In some instances, this led to the chameleon accidentally wrapping its tail around a hind leg, still holding on to the perch from which they were departing and causing them to fall. In the wild, falling from a perch could have significant fitness and energetic consequences as it increases visibility and forces animals to climb the tree to return to their perches. It might be that animals bred and kept in captivity are bolder and take more risks. Consequently, this behavior might be seen less often in wild individuals. In the wild, where natural perches are likely rougher and more irregular in diameter and surface texture than the ones used for this experiment, there is likely no need to develop different strategies to deal with smoother substrates. While they are highly capable of crossing distances greater than their own shoulder-hip length, the extent of their capabilities relies on the coarseness and diameter of the substrate. Very smooth substrates created difficulties for the chameleons and appeared to prevent them from generating enough hand and feet grip strength to hold on to the perches. Consequently, very few successful crossings were observed with PVC as a substrate (only a few successes were made at 0.5 and 1 time the shoulder-hip distance). When on VC perches, the animals were more likely to fall off from the perch before the trial even started, adding to the argument that chameleons rely on the friction of the substrate as well as on their own grasping strength.

### Sex related size differences and tail use preference

Our results showed that males were more successful in crossing greater distances than females (Figs. 2, 3). As the distances were adjusted to the average length per sex, it seems that there is a difference in performance difference irrespective of size. Overall, males more often did not use their tails when crossing, whereas females only occasionally crossed the shortest distance without using their tails. From distances equal to shoulder-hip length onwards males do use their tail more often than not. *Chamaeleo calyptratus* is a sexually dimorphic species with the males being greater in size in general, having larger hands and feet and longer tails. It might be expected that males would perform less well due to their average larger size as muscle force increase less fast than body mass (Meyers et al., 2002; Herrel et al., 2006a; Herrel et al., 2006b). While muscle force typically scales with negative allometry relative to mass, the males are much larger than females so we can expect that they would be stronger both in absolute. Yet positive allometry is often seen in functional traits with force increasing faster than predicted by geometric scaling (Herrel and Gibb, 2006). Moreover, males are likely under strong selection to perform well. When males encounter each other, they will often engage in fights during which they will attempt to toss their rival from the branch onto the ground (Measey et al., 2009). Sexual selection for males with a higher gripping performance could consequently explain why male chameleons perform better for their size. Another factor that could be considered is the boldness of animals. This may explain why males tended to attempt crossing greater distances, whereas females did not. Boldness as a personality trait has been described in other squamates, such as for the Namibian rock agama, *Agama planiceps* (Carter et al., 2012). Exploring personality and its links to morphology, performance, and behaviors like gap-bridging would be a worthwhile avenue for future research.

## Data accessibility

Data are available in the supplementary materials, video data can be acquired upon request.

## Acknowledgements

The authors would like to thank Jens De Meyer, Aurélien Lowie, Simon Baeckens, Robert Paauwe, and the anonymous reviewers for their feedback; Barbara De Kegel and Eva Vangenechten for their assistance in the animal caretaking and Alexis Dollion for providing us with the chameloens. This study was funded by the Fonds Wetenschappelijk Onderzoek [#3G006716] and a Tournesol mobility grant.

Version 3 of this preprint has been peer-reviewed and recommended by Peer Community In Zoology (https://doi.org/10.24072/pci.zool.100005)

## Conflict of interest disclosure

The authors of this preprint declare that they have no financial conflict of interest with the content of this article.

Prof. dr. Dominique Adriaens is one of the *PCI Zool* recommenders.

## References

Ali, S.M. (1947). Studies on the anatomy of the tail in sauria and rhynchocephalia. Proceedings of the Indiana Academy of Science 28: 151–165

Andrews, R.M. (2008) Lizards in the slow lane: Thermal Biology of chameleons. Journal of Thermal Biology 33: 57–61.

Bergmann, P.J., Lessard, S. & Russel, A.P. (2003) Tail growth in Chamaeleo dilepis (Sauria: Chamaeleonidae): functional implications of segmental patterns. Journal of Zoology 261: 417–425

Bickel, R. and Losos, J.B. (2002). Patterns of morphological variation and correlates of habitat use in Chameleons. Biological Journal of the Linnean Society 76: 91–103.

Byrnes G. & Jayne, B.C. (2012) The effects of three-dimensional gap orientation on bridging performance and behavior of brown tree snakes (Boiga irregularis). Journal of Experimental Biology 215: 2611–2620.

Carter, A.J., Heinsohn, R.I., Goldizen, A.W. & Biro, P.A. (2012) Boldness, trappability and sampling bias in wild lizards. Animal Behavior 83: 1051–1058.

Cartmill, M. (1985). Climbing. In Functional Vertebrate Morphology (ed. M. Hildebrand, D. M. Bramble, K. F. Liem and D. B. Wake), pp. 73–88. Cambridge: Belknap Press.

da Silva, J., A. Herrel, G.J. Measey, B. Vanhooydonck & Tolley, K.A. (2014) Linking microhabitat structure, morphology and locomotor performance traits in a recent radiation of dwarf chameleons (Bradypodion). Functional Ecology 28: 702–713

Fischer, S.M., Krause, C. & Lilje, K.E. (2010) Evolution of chameleon locomotion, or how to become arboreal as a reptile. Zoology 113: 67–74.

Gans, C. (1967). The chameleon. Natural History 76: 52–59.

Graham, M. & Socha, J.J. (2020) Going the distance: The biomechanics of gap-crossing behaviors. Journal of Experimental Zoology A: Ecological and Integrative Physiology 333: 60–73.

Herrel, A. & Gibb, A.C. (2006a) Ontogeny of performance in vertebrates. Physiological and Biochemical Zoology 79: 1–6

Herrel, A. & O’Reilly, J.C. (2006b) Ontogenetic scaling of bite force in lizards and turtles. Physiological and Biochemical Zoology 79: 31–42

Herrel, A., Measey, G.J., Vanhooydonck, B. & Tolley, K.A. (2011). Functional consequences of morphological differentiation between populations of the Cape Dwarf Chameleon (Bradypodion pumilum). Biological Journal of the Linnean Society 104: 692–700

Herrel, A., Measey, G.J., Vanhooydonck, B. & Tolley, K.A. (2012) Got it clipped? The effect of tail clipping on tail gripping performance in chameleons. Journal of Herpetology 46: 91–93

Herrel, A., Tolley, K.A., Measey, G.J., da Silva, J.M., Potgieter, D.F., Boller, E., Boistel, R. & Vanhooydonck, B. (2012). Slow but tenacious: an analysis of running and gripping performance in chameleons. Journal of Experimental Biology 216: 1025–1030

Hoefer, K.M. & Jayne, B.C. (2013) Three-dimensional locations of destinations have species-dependent effects on the choice of paths and the gap-bridging performance of arboreal snakes. Journal of Experimental Zoology A: Ecological Genetics and Physiology 319: 124–37

Khannoon, E.R., Endlein, T., Russel, A.P., Autumn, K. (2014) Experimental evidence for friction-enhancing integumentary modifications of chameleons and associated functional and evolutionary implications. Proceedings of the Royal Society B. 281(1775).

Luger, A.M., Ollevier, A., De Kegel, B., Herrel, A. and Adriaens, D. (2020). Is variation in tail vertebral morphology linked to habitat use in chameleons? Journal of Morphology 281: 229–239

Measey, G.J., Hopkins, K.P. & Tolley, K.A. (2009). Morphology, ornaments and performance in two chameleon ectomorphs: is the casque bigger than the bite? Zoology 112: 217–226

Meyers, J.J., A. Herrel & J. Birch (2002) Scaling of morphology, bite force, and feeding kinematics in an iguanian and a scleroglossan lizard. In: Topics in Functional and Ecological Vertebrate Morphology (P. Aerts, K. D’aout, A. Herrel and R. Van Damme, Eds.). Shaker Publishing, Maastricht. pp. 47–62

Peterson, J.A (1984) The locomotion of Chamaeleo (Reptilia: Sauria) with particular reference to the forelimb. Journal of Zoology 202:1–42

Spinner, M., Westhoff, G., Gorb, S.N. (2014) Subdigital setae of chameleon feet: friction-enhancing microstructures for a wide range of substrate roughness. Scientific Reports. 4(5481)

Thorpe, S. K. S., Holder, R. and Crompton, R. H. (2009) Orangutans employ unique strategies to control branch flexibility. Proceedings of the National Academy of Sciences 106: 12646–12651

Zippel, K.C. & Glor, R.E. (1999). On caudal prehensility and phylogenetic constraint in lizards: the influence of ancestral anatomy on function in Corucia and Furcifer. Journal of Morphology 239: 143–155

