## Supplementary material for "Do substrate roughness and gap distance impact gap-bridging strategies in arboreal chameleons": Suppl. table 2

Suppl. Table 2: Experimental data giving the total number of times each set up was successfully crossed by the chameleon *Chamaeleo calyptratus* with or without using their tail (in absolute numbers and percentages). Each set-up is organized according to materials (PVC, sandpaper and wood), perch thickness (narrow or broad), gap distance relative to the shoulder-hip length of each sex, and sex.

| **Material** | **Sex** | **Thickness** | **Gap distance** | **number tail used** | **number tail not used** | **Tail used %** | **Tail not used %** |
| --- | --- | --- | --- | --- | --- | --- | --- |
| PVC | F | 25 mm | 0.5 | 0.0 | 1 | 0.0 | 100 |
|  | F |  | 1 |  |  |  |  |
|  | F |  | 1.25 |  |  |  |  |
|  | F |  | 1.5 |  |  |  |  |
|  | F |  | 1.75 |  |  |  |  |
|  | F |  | 2 |  |  |  |  |
|  | F | 9 mm | 0.5 | 0.0 | 1 | 0.0 | 100 |
|  | F |  | 1 |  |  |  |  |
|  | F |  | 1.25 |  |  |  |  |
|  | F |  | 1.5 |  |  |  |  |
|  | F |  | 1.75 |  |  |  |  |
|  | F |  | 2 |  |  |  |  |
|  | M | 25 mm | 0.5 | 0.0 | 3 | 0.0 | 100 |
|  | M |  | 1 |  |  |  |  |
|  | M |  | 1.25 |  |  |  |  |
|  | M |  | 1.5 |  |  |  |  |
|  | M |  | 1.75 |  |  |  |  |
|  | M |  | 2 |  |  |  |  |
|  | M | 9 mm | 0.5 | 2 | 3 | 40 | 60 |
|  | M |  | 1 | 3 | 0.0 | 100 | 0.0 |
|  | M |  | 1.25 |  |  |  |  |
|  | M |  | 1.5 |  |  |  |  |
|  | M |  | 1.75 |  |  |  |  |
|  | M |  | 2 |  |  |  |  |
| Sandpaper | F | 25 mm | 0.5 | 3 | 6 | 33.33 | 66.67 |
|  | F |  | 1 | 8 | 0.0 | 100 | 0.0 |
|  | F |  | 1.25 | 6 | 0.0 | 100 | 0.0 |
|  | F |  | 1.5 | 6 | 0.0 | 100 | 0.0 |
|  | F |  | 1.75 |  |  |  |  |
|  | F |  | 2 |  |  |  |  |
|  | F | 9 mm | 0.5 | 1 | 7 | 12.5 | 87.5 |
|  | F |  | 1 | 8 | 0.0 | 100 | 0.0 |
|  | F |  | 1.25 | 8 | 0.0 | 100 | 0.0 |
|  | F |  | 1.5 | 8 | 0.0 | 100 | 0.0 |
|  | F |  | 1.75 | 3 | 0.0 | 100 | 0.0 |
|  | F |  | 2 |  |  |  |  |
|  | M | 25 mm | 0.5 | 4 | 6 | 40 | 60 |
|  | M |  | 1 | 8 | 5 | 80 | 20 |
|  | M |  | 1.25 | 9 | 1 | 90 | 10 |
|  | M |  | 1.5 | 10 | 0.0 | 100 | 0.0 |
|  | M |  | 1.75 | 3 | 0.0 | 100 | 0.0 |
|  | M |  | 2 |  |  |  |  |
|  | M | 9 mm | 0.5 | 2 | 6 | 25 | 75 |
|  | M |  | 1 | 5 | 3 | 62.5 | 37.5 |
|  | M |  | 1.25 | 7 | 1 | 87.5 | 12.5 |
|  | M |  | 1.5 | 5 | 0.0 | 100 | 0.0 |
|  | M |  | 1.75 | 3 | 0.0 | 100 | 0.0 |
|  | M |  | 2 |  |  |  |  |
| Wood | F | 25 mm | 0.5 | 9 | 5 | 64.29 | 35.71 |
|  | F |  | 1 | 10 | 0.0 | 100 | 0.0 |
|  | F |  | 1.25 | 3 | 0.0 | 100 | 0.0 |
|  | F |  | 1.5 |  |  |  |  |
|  | F |  | 1.75 |  |  |  |  |
|  | F |  | 2 |  |  |  |  |
|  | F | 9 mm | 0.5 | 3 | 13 | 18.75 | 81.25 |
|  | F |  | 1 | 17 | 0.0 | 100 | 0.0 |
|  | F |  | 1.25 | 5 | 0.0 | 100 | 0.0 |
|  | F |  | 1.5 | 10 | 0.0 | 100 | 0.0 |
|  | F |  | 1.75 |  |  |  |  |
|  | F |  | 2 |  |  |  |  |
| Wood | M | 25 mm | 0.5 | 8 | 5 | 61.54 | 38.46 |
|  | M |  | 1 | 10 | 2 | 83.33 | 16.67 |
|  | M |  | 1.25 | 8 | 0.0 | 100 | 0.0 |
|  | M |  | 1.5 | 2 | 0.0 | 100 | 0.0 |
|  | M |  | 1.75 |  |  |  |  |
|  | M |  | 2 |  |  |  |  |
|  | M | 9 mm | 0.5 | 9 | 13 | 40.9 | 59.09 |
|  | M |  | 1 | 17 | 5 | 77.27 | 22.73 |
|  | M |  | 1.25 | 10 | 0.0 | 100 | 0.0 |
|  | M |  | 1.5 | 10 | 1 | 90.9 | 9.09 |
|  | M |  | 1.75 |  |  |  |  |
|  | M |  | 2 |  |  |  |  |
