## Supplementary material for "Do substrate roughness and gap distance impact gap-bridging strategies in arboreal chameleons": Suppl. table 1

Suppl. Table 1: Measurements of the chameleons used. Weight is given in gram, length in millimeter. Males are listed as M1-5, females F1-4.

|  | **M1** | **M2** | **M3** | **M4** | **M5** | **F1** | **F2** | **F3** | **F4** |
| --- | --- | --- | --- | --- | --- | --- | --- | --- | --- |
| **Weight** | 193 | 197 | 147 | 187 | 158 | 135 | 197 | 180 | 137 |
| **Tail length (cloaca – tail tip)** | 220 | 214 | 212 | 206 | 209 | 156 | 160 | 156 | 155 |
| **Snout-vent length** | 170 | 175 | 165 | 180 | 170 | 140 | 155 | 145 | 145 |
| **Shoulder-hip length** | 133 | 131 | 128 | 133 | 125 | 112 | 110 | 108 | 110 |
